# Neuropeptide Signaling Promotes Stem-Cell Driven Regeneration in Planarians

**DOI:** 10.64898/2026.10.10.758126

**Authors:** Christina E. Endara-Arnold, Jennifer E. Jenkins, Andres E. Villalobos, Rachel H. Roberts-Galbraith

**Affiliations:** University of Georgia, Neuroscience Program, 120 Cedar St., Athens, GA, 30602, USA; University of Georgia, Cellular Biology Department, Current Address: Duke University, Cellular Biology Department, 307 Research Drive Durham, NC 27710; University of Georgia, College of Engineering, Current Address: Northwestern University, 355 E Erie St, Chicago, Illinois 26N

## Abstract

Neurons and neuroendocrine cells produce neuropeptides, which act as long-range signals to modulate organismal state and physiological function. In a few cases, neuropeptides have been thought to promote regeneration, but understanding this connection has been difficult due to the relatively poor regenerative capacity of most traditional model organisms. To test whether neuropeptides drive robust regeneration, we used the highly regenerative asexual planarian flatworm as a model. We began by investigating enzymes that cleave and process pre-propeptides to produce bioactive neuropeptides. We characterized 12 putative neuropeptide-processing enzymes and identified three enzymes, prohormone convertase 2 (pc2), carboxypeptidase E (CPE), and peptidylglycine alpha-hydroxylating monooxygenase (PHM-1) that regulate regeneration through their effects on stem cell differentiation and maintenance. We then investigated 51 of the 52 known neuropeptide-encoding genes in planarians, finding that 16 of these genes promote efficient regeneration and/or stem cell function. Our work indicates that diverse neuropeptides regulate endogenous pluripotent stem cells during homeostasis and in robust regeneration. Our data supports a key role for neuropeptide signaling in planarians to promote a pro-regenerative state, which could be valuable to mimic in regenerative therapies.

**Significance Statement:** Planarians possess pluripotent stem cells that give rise to all cell types in regeneration and homeostasis. Importantly, the signaling pathways that control stem cell behaviors in planarians, particularly after injury, are poorly understood. We here test the hypothesis that the planarian nervous system is the source of key signals that allow proper planarian regeneration to occur. Our work further reveals that the planarian nervous system promotes signaling for tissue renewal and stem cell control through a diverse group of neuropeptides.

## Introduction

After injury, regeneration allows cells, tissues, or organs to be replaced to restore organismal health and function. In many organisms, the nervous system itself influences regenerative state. Famously, in newts and axolotls, Schwann cells produce anterior gradient protein to aid in limb regeneration (Kumar, 2012). Similarly, innervation of the fin is critical for regenerative success in Zebrafish (Simões et al., 2014). On the other hand, neurons are largely dispensable for regeneration in *Hydra* and in the early larval *Xenopus* limb, but even in these cases neurons facilitate an optimal signaling environment to expedite regeneration (Filoni et al., 1999; Marcum & Campbell, 1978; Pirotte et al., 2016). Taken together, neurons play diverse roles in regeneration across animal phyla, but the specific mechanisms through which neurons influence regeneration are often uncharacterized, and the universality of neuronal control of regeneration is not yet clear, particularly in systems capable of whole-body regeneration.

Freshwater flatworms called planarians have emerged as an outstanding model for discovery and elucidation of cell signaling pathways that impact robust whole-body regeneration. Planarians regenerate through the action of a population of pluripotent stem cells called neoblasts (Wagner et al., 2011). These stem cells give rise to all adult cell types in the planarian and replace missing tissue after injury (Wagner et al., 2011). Hints that the planarian nervous system promotes the function of stem cells, exist. For example, sub-lethal irradiation of planarians showed surviving stem cells preferentially nestled near the nervous system (Rossi et al., 2018). Some signaling molecules expressed in planarian neurons also modulate stem cell function, including Neuregulin-7, which modulates the symmetry of stem cell division (Lei et al., 2016). Still, how and whether the planarian nervous system influences stem cells during the process of regeneration is still unclear.

One feature of the nervous systems in animals is production of neurotransmitters and neuropeptides that activate neuronal signaling through synapses and in some cases through long-range signaling and neuromodulation (Nadim & Bucher, 2014; van den Pol, 2012). Perturbation of planarian prohormone convertase 2 (PC2), a neuropeptide processing enzyme, was previously shown to cause paralysis and to promote formation of unpigmented new tissue called the blastema (P. W. Reddien, A. L. Bermange, et al., 2005). As seen in other animals, planarian PC2 is an enzyme that cleaves prepropeptides at dibasic residues, which allow neuropeptides to become activated (Fig. 1A, (Van Bael et al., 2018)). After prohormone convertases cleave prepropeptides (Sharker et al., 2020), carboxypeptidases remove dibasic sequences (Song & Fricker, 1995) and peptidylglycine-α-amidating monooxygenase (PAM) enzymes catalyze their amidation (Eipper et al., 1992), allowing full maturation and bioactivity of neuropeptides (Fig. 1A). Mature neuropeptides are secreted from dense core vesicles as neuromodulatory, hormonal, and/or endocrine signals (Kirchner et al., 2023) that regulate neurodevelopment, neurotransmission, behavior, energy homeostasis, and other physiological properties of animal species (Aponte et al., 2011; Cohen et al., 2009; Russo, 2017; Sakurai & Mieda, 2011). Intriguingly, neuropeptides secreted by neurons, glia, and other endocrine tissues (Hur et al., 2010) play emerging roles in regeneration and in regulation of stem cells in other species (Park et al., 2015; Singh et al., 2017; Xu et al., 2023). However, PC2 does process non-neuropeptide substrates in other animals (Ohagi et al., 1992; Rouillé et al., 1994), so it is not yet clear whether planarian PC2 promotes movement and regeneration through maturation of neuropeptides or other substrates.

**Figure 1.**
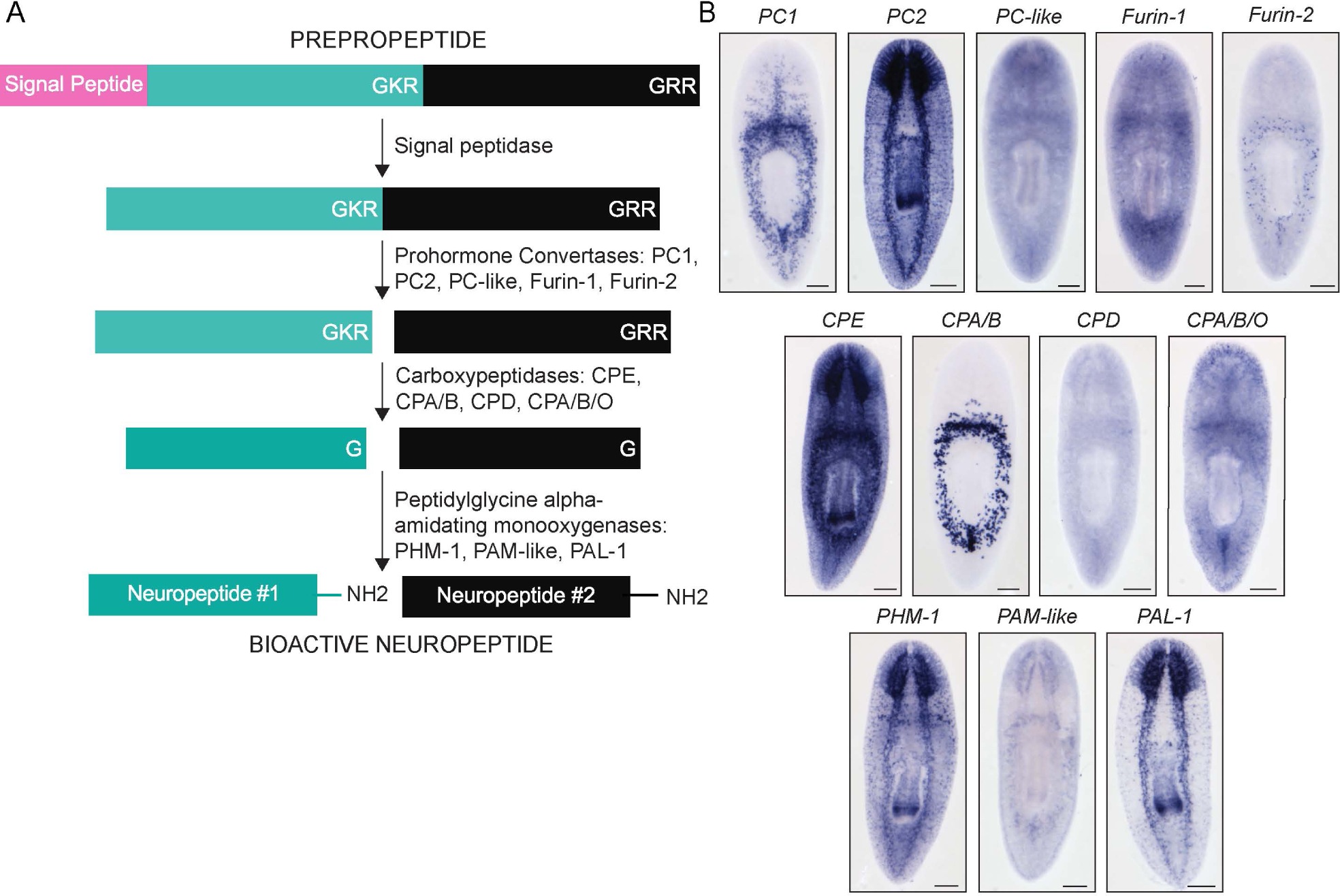
Identification of planarian neuropeptide processing enzyme-encoding genes. (A) Diagram of the neuropeptide biosynthesis pathway adapted from (Van Bael et al., 2018). Each step is indicated with an arrow and annotated with the implicated neuropeptide processors. Dibasic residues are represented by K (lysine) and R (arginine). Glycine is represented by the letter G. (B) Untreated *in situ* hybridization (ISH) experiments show expression of twelve planarian neuropeptide processing enzyme-encoding genes. Scale bars = 200 µm (N=8-12 worms).

Planarian PC2 has been definitively linked to neuropeptide processing and at least 52 planarian neuropeptides have been identified (Collins et al., 2010; Ong et al., 2016). Each prepropeptide-encoding gene is expressed in unique patterns through the central nervous system, peripheral nervous system, eyes, and pharynx (Collins et al., 2010; Ong et al., 2016). A handful of neuropeptides have been further characterized, including neuropeptide Y-8 (NPY-8) which is essential for development and maintenance of testes and production of sperm in sexual planarians (Collins et al., 2010). In the planarian species *Dugesia japonica*, pharyngeal extension behavior requires five neuropeptide prohormones (Ishii, 1965; Macrae, 1963; Shimoyama et al., 2016; Wulzen, 1917). One neuropeptide, insulin-like peptide-1,(ILP-1) promotes stem cell proliferation and body growth (Miller & Newmark, 2012). Other neuropeptides or peptidigeric treatments in planarians have been implicated in roaming behavior, water regulation, cell proliferation, cell differentiation, pharynx regeneration, and sensory filtering (Baguna et al., 1989; Bautz & Schilt, 1986; Bray et al., 2024; Hori, 1997; Hori & Kishida, 2003; Kobayashi et al.; Kreshchenko, 2008; Kreshchenko et al., 2008; Saló & Baguñà, 1986; Wyss et al., 2024). But the role of most neuropeptides and thus peptidergic neurons are unclear, particularly in the context of stem cell modulation and regeneration.

To uncover whether the planarian nervous system promotes regeneration, we began by characterizing functions for the full complement of planarian neuropeptide processing enzymes. We examined each gene’s expression pattern and functions, finding that planarian *PC2*, *CPE*, and *peptidylglycine alpha-hydroxylating monooxygenase-1* (*PHM-1*) are co-expressed and are essential for regeneration, stem cell maintenance, stem cell differentiation, and animal behavior. Our results support the hypothesis that planarian PC2 and its associated partners promote neuropeptide processing to promote regeneration and stem cell function. We further tested this hypothesis by identifying sixteen prepropeptide-encoding genes that are individually required for optimal brain regeneration. Importantly, five of these neuropeptides influenced the ability of stem cells to proliferate adequately after injury, while others play stem cell-independent roles. Collectively, our data demonstrate that the planarian nervous system is critical for whole-body regeneration, through production of neuropeptides that play diverse, essential roles. We speculate that neuropeptides in planarians and potentially other animals signal to create a pro-regenerative physiological state.

## Results

### Planarian neuropeptide processing enzyme homologs

In most animals, neuropeptides become bioactive upon cleavage by processing enzymes (Fig. 1A). We predicted that if neuropeptides play a vital role in planarian regeneration, one straightforward way to detect this role would be through perturbation of the neuropeptide processors important for their synthesis. We identified twelve genes predicted to encode neuropeptide processors and verified their assignments and homology through mutual BLAST and phylogenetic analysis (Supp. Table 1, Supp. Fig. 1, 2). Processers included: two prohormone convertases (*PC1* and *PC2*); one *prohormone convertase-like* gene (PC-like); two *Furin* homologs (Furin-1 and Furin-2); and four carboxypeptidases (*CPE, CPA/B, CPD*, and *CPA/B/O*). We also found homologs for peptidylglycine alpha-amidating monooxygenase (*PAM-like*) and the two catalytic domains of PAM which are Peptidyl-α-amidating lyase (*PAL-1*) and Peptidylglycine-α-hydroxylating monooxygenase (*PHM-1*) (Asada et al., 2005). We cloned each gene and determined where each putative neuropeptide processor-encoding gene was expressed in the planarian body using *in situ* hybridization (ISH). The majority of processor- encoding genes were expressed in the central nervous system and peripheral nervous system (Fig. 1B). However, *PC1* expression was highest in or near the intestine, while *CPA/B* and *Furin- 2* expression were most obvious in peripharyngeal puncta. Several homologs were also expressed in the pharynx, including *PC2*, *CPE*, *PHM-1*, and *PAL-1*. Our ISH results largely confirm data from single-cell transcriptomic data (Fincher et al., 2018) (Supp. Table 2).

### *PC2* is essential for regeneration, stem cell function, and planarian locomotion

Perturbation of *PC2* was previously shown to cause paralysis, small blastemas, and lysis (P. W. Reddien, A. L. Bermange, et al., 2005). We repeated RNA interference (RNAi) targeting *PC2* and observed immobility and ventral curling after only one or two dsRNA feedings as previously reported (Supp. Fig. 3A-B). We also observed pharynx ejection in *PC2(RNAi)* worms (Fig. 2A-B) as well as subsequent lesions and eventual lysis (Fig. 2C). Taken together, our results confirm that PC2 is needed for proper locomotion and planarian viability (Supp. Fig. 3C).

**Figure 2.**
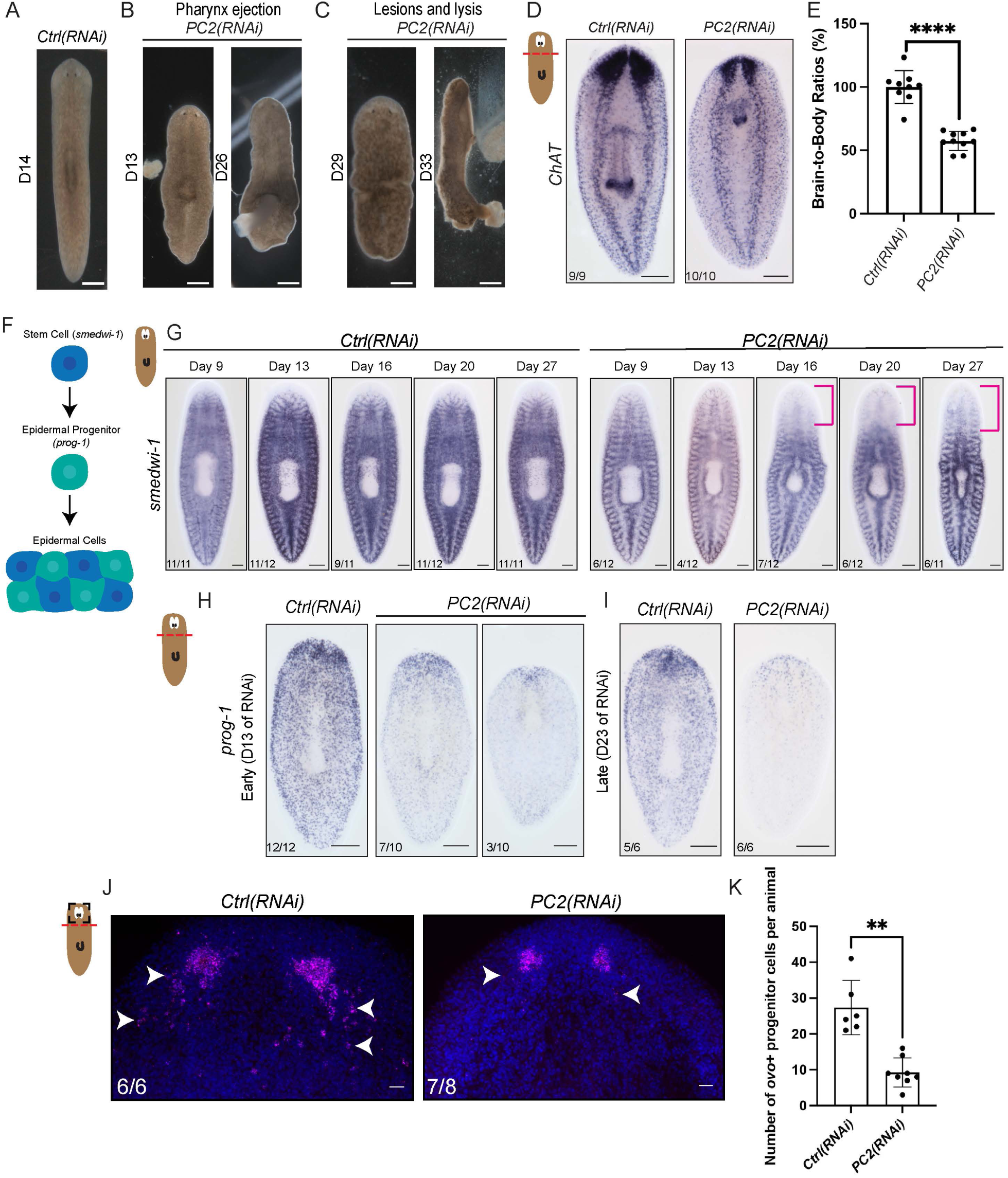
Prohormone convertase 2 (*PC2*) is vital for survival, regeneration, and stem cell differentiation. (A) Image of a representative negative control worm that was fed dsRNA matching *green fluorescent protein* (*gfp*) two times. (B) Representative image of worms fed *PC2* dsRNA displaying ejection and detachment of the feeding organ (pharynx). (C) *PC2(RNAi*) worms show tissue integrity issues, including lesions and lysis. The RNAi paradigm consisted of two dsRNA feedings 3 days apart, with “D#” indicating the number of days lapsed since the first RNAi feeding. (D) Visualization of the central nervous system with *choline acetyltransferase* (*ChAT)* 7 days after head amputation. (E) Bar graph displaying brain-to-body ratios produced by tracing the brain and body in triplicate (N= 9-10 worms) and normalizing such that controls were set to 100%. Unpaired Welch’s test with Welch’s correction, graph showing mean with standard deviation. (F) Diagram displaying the pathway from a pluripotent stem cell to an epidermal progenitor (*prog-1^+^*) which will become mature epidermal cells (Cheng et al., 2018). (G) *in situ* hybridization (ISH) with stem cell marker, *smedwi-1,* in intact worms after 3 dsRNA feeds. Magenta brackets indicate areas of reduced *smedwi-1* mRNA staining. (H) Regenerating RNAi animals stained with *prog-1* (epidermal progenitor marker) 5 days post-amputation at an early time point (13 days of RNAi) (I) and a late time point (23 days of RNAi). (J) Head regenerated animals stained with *ovo* at 5 days post-amputation (N= 6-8 worms). White arrowheads indicate *ovo^+^*eye progenitors. (K) Bar graph displaying the number of *ovo^+^* cells in head regenerates. Unpaired t-test with Welch’s correction showing the mean with standard deviation. **** p value ≤ 0.001; ** p value = 0.0010. Scale bars A-C = 500 µm; D-I = 200 µm; J = 20 µm.

We next investigated the degree to which planarian *PC2* promotes robust tissue regeneration. Previously, Reddien and colleagues had shown in a large-scale screen that *PC2* knockdown leads to reduced head blastema size (Reddien, Bermange, et al., 2005). We confirmed that *PC2*(RNAi) leads to smaller head *and* tail blastemas (Supp. Fig. 3D-E) and causes an even more marked impact on brain regeneration, which we visualized by staining with *choline acetyltransferase* (*ChAT*) and quantified through brain area normalized to body size (Fig. 2D-E). We conclude that multiple forms of regeneration are significantly stunted in the absence of *PC2*.

Pharynx ejection and epidermal lesions have previously been seen after perturbation of pharyngeal (Adler et al., 2014) or epidermal progenitors respectively (van Wolfswinkel et al., 2014; Wagner et al., 2012). We therefore wondered if PC2 might be important for planarian stem cell maintenance or function. We first investigated whether intact *PC2(RNAi)* animals lose their stem cells over time, following RNAi with ISH to observe *smedwi-1*, a marker for planarian stem cells (P. W. Reddien, N. J. Oviedo, et al., 2005). Even before lesions or lysis phenotypes arose, we saw a consistent decrease in *smedwi-1* staining as early as day 9 of RNAi (Fig. 2F- G). We also saw that *smedwi-1* transcript was more significantly reduced in *PC2(RNAi)* animals regenerating their heads, which was confirmed through RT-qPCR (Supp. Fig. 3F-H). Additionally, we saw that *bruli*, another stem cell marker, (Guo et al., 2006) was also reduced in *PC2(RNAi)* regenerating worms (Supp. Fig 3H). Our results show that *PC2* is required for stem cell maintenance in both regeneration and non-injury settings.

We next tested whether stem cell differentiation is affected by *PC2* perturbation. We examined epidermal progenitors using the marker *prog-1* (Eisenhoffer et al., 2008) and noted a dramatic reduction in *prog-1^+^* cells after *PC2(RNAi)* at early (day 13) and late (day 23) timepoints (Fig. 2H-I). These results indicate that *PC2* expression supports production and/or maintenance of differentiating epidermal lineage progenitors. We then investigated whether progenitor depletion occurs for other lineages. The transcription factor-encoding gene *ovo* is expressed in the eye and in eye progenitors, that appear in visible trails posterior to the eye (Lapan & Reddien, 2012). In regenerating *PC2(RNAi)* worms, we saw fewer eye progenitors (Fig. 2J-K). Taken together, we conclude that stem cell differentiation is broadly reduced in the absence of *PC2*.

### CPE and PHM-1 cooperate with PC2 to promote regeneration and stem cell function

Planarian *PC2* could promote proper regeneration, viability, and stem cell function through neuropeptide processing, but there are also examples of prohormone convertases and furins in other species cleaving proteins that are unrelated to neuropeptides, including extracellular matrix proteins (e.g. glypican, (De Cat et al., 2003), signaling molecules (e.g. TGFβ, (Dubois et al., 1995), and receptors (e.g. insulin receptor, (Baker et al., 1993). To bolster our model that planarian *PC2* operates through neuropeptides, we next sought to determine whether other planarian neuropeptide processors played similar roles in regeneration and stem cell biology. We reasoned that if perturbation of other neuropeptide processors recapitulated *PC2*(RNAi) phenotypes, neuropeptides would be the most relevant PC2 target.

We began by investigating roles for each neuropeptide-processing enzyme in central nervous system regeneration, given that this assay is a sensitive readout of regenerative success. We determined that *CPE(RNAi*) and *PHM-1*(*RNAi)* animals displayed statistically smaller brain size after head amputation and regeneration compared to control animals (Fig. 3A-B), while perturbation of other putative processors caused no significant impact on regeneration (Supp. Fig. 4A-B). Interestingly, we also noted behavioral and tissue maintenance phenotypes in *CPE(RNAi)* animals, including lesions (Fig. 3C). We also observed that *CPE(RNAi)* worms also impacted survival in a non-amputation condition, but we did not see the same lethality in *PHM-1(RNAi)* animals (Supp. Fig 4C). Then, we *CPE(RNAi)* animals displayed slow movement and these animals took significantly longer to right themselves when placed on their dorsal side, both pre- and post-amputation, reflecting movement difficulties (Supp Figs. 4D- E). Our results suggested that *CPE* and *PHM-1* encode the most likely partners of PC2 in promoting regeneration and that *CPE* likely cooperates with *PC2* in regulation of planarian behavior.

**Figure 3.**
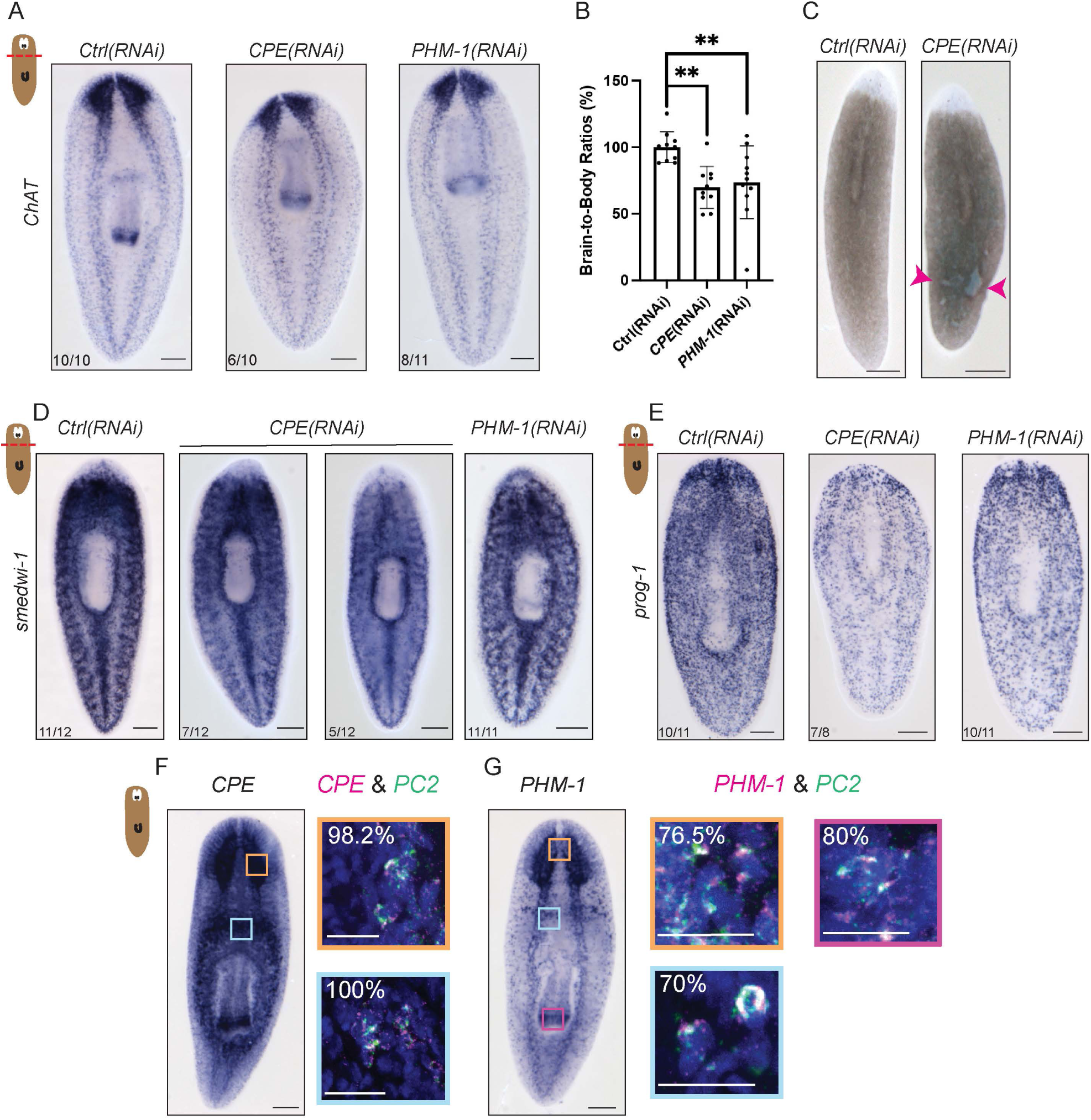
*CPE* and *PHM-1 cooperate with PC2* in regeneration. (A) Representative images showing brain regeneration after head amputation in *CPE(RNAi)* and *PHM-1(RNAi)* animals. The RNAi paradigm consisted of 5 µg dsRNA feedings every 3-4 days, head amputation after 7 days, and 7 days for head regeneration. (B) Quantification of A with analysis and One-way ANOVA with Dunnett’s multiple comparisons (N=10-11 worms). Graph showing means with standard deviation with controls normalized to 100% to ease comparisons. (C) *CPE(RNAi)* animals after five RNAi feeds and amputation displaying lesions as shown by magenta arrowheads. (D-E) Representative regenerating worms stained with *smedwi-1* (stem cell marker) (Peter W. Reddien et al., 2005) or *prog-1* (epidermal progenitor marker) (Cheng et al., 2018), after neuropeptide processor knockdowns. (N=8-12 worms) (F) Double fluorescent *in situ* hybridization showing *Carboxypeptidase E* (*CPE*) and *PC2* coexpression in the brain (orange square) and peripheral nervous system (light blue square). Percentages of *CPE^+^* cells that are *PC2^+^* are indicated in the top left corner of each inset image. (G) Double fluorescent *in situ* hybridization displaying *PHM^+^*cells coexpressing *PC2* in the brain (orange square), peripheral nervous system (light blue square), and pharynx (pink square). Percentages of *PHM-1*^+^ cells coexpressing *PC2^+^* are indicated in the top left corner of each image. (N= 6 worms; n= 180-300 cells quantified per body region) **p value = 0.0035. Scale bars = 200 µm; except inset scale bars in F-G = 50 µm.

We predicted that *CPE* and *PHM-1*, like *PC2*, impact regeneration through effects on stem cells. Through ISH and RT-qPCR, we confirmed that *smedwi-1* and *prog-1* transcripts were reduced in *CPE(RNAi)* and *PHM-1(RNAi)* worms, with the change in staining most obvious near the wound site (Fig. 3D-E and Supp. Fig. 4F-G). We also investigated the roles for both genes in homeostasis and saw that *CPE(RNAi)* and *PHM-1*(RNAi) animals had less *prog-1*^+^ staining even in the absence of injury (Supp. Fig. 4H). We also noted that homeostatic *CPE(RNAi)* worms had statistically smaller brain size compared to control worms, suggesting failure to maintain neural tissues (Supp Fig. 4I-J). Our data indicate that knockdown of *CPE* or *PHM-1* partially phenocopies loss of *PC2*. Overall, the phenotypes of *CPE*(RNAi) and *PHM- 1*(RNAi) animals were less severe than *PC2*(RNAi) animals, potentially reflecting partial bioactivity of incompletely processed neuropeptides previously noted in other organisms (Liu et al., 1995; Pan et al., 2006).

Finally, *CPE* and *PHM-1* must be co-expressed with *PC2* in order for the enzymes they encode to work together to process common neuropeptide substrates. To confirm coexpression, we performed double fluorescent *in situ* hybridization (dFISH) experiments. *CPE* was highly co-expressed with *PC2* in the brain (98.2% of *CPE*^+^ cells are *PC2*^+^) and the peripheral nervous system (100% coexpression) (Fig. 3F). Over 70% of PHM-1^+^ cells also expressed *PC2* in the brain, “neck”, and periphery of the planarian (Figure 3G). In contrast, the other neuropeptide processor-encoding genes were expressed in patterns that overlapped less frequently with *PC2*, though the remaining overlap could indicate some cooperation (Supp Figs. 5A-R). Shared functions and colocalization of *CPE and PHM-1* with *PC2* supports a model in which these processors work together to create bioactive neuropeptides important for regeneration, stem cell function, and behavior.

### Neuropeptide-encoding genes implicated in regeneration and stem cell function

Given our support for the hypothesis that neuropeptides promote planarian regeneration, we ultimately tested the function of all the known neuropeptides expressed in asexual *Schmidtea mediterranea,* omitting *npy-8* since its expression is specific to sexual planarians of the same species (Collins et al., 2010; Ong et al., 2016). We knocked down each neuropeptide-encoding gene and measured regenerated brain size, which was normalized to body size. RNAi of sixteen neuropeptide-encoding genes led to statistically smaller brains after regeneration, though some of these phenotypes were subtle (Fig. 4A-B). To test whether regenerative roles are specific to the brain or general to all body parts, we also looked at the ability of animals to regenerate a pharynx, the feeding organ. We determined that one pharynx-expressed neuropeptide-encoding gene*, pyrokinin prohormone like-1*, *ppl-1*, was essential for regenerating pharynges of typical size relative to body size (Fig. 4C-D). Because neuropeptides can work cooperatively, we also tested whether simultaneous knockdown of the top six neuropeptides, *spp-14, spp-15*, *spp-16*, *spp-18*, *spp-20*, and *ppl-1,* caused a more severe brain regeneration phenotype. Knocking down these six neuropeptides at the same time did not lead to a more robust phenotype, but RNAi of so many genes simultaneously might have impaired knockdown efficiency (Supp. Fig. 6A-D).

**Figure 4.**
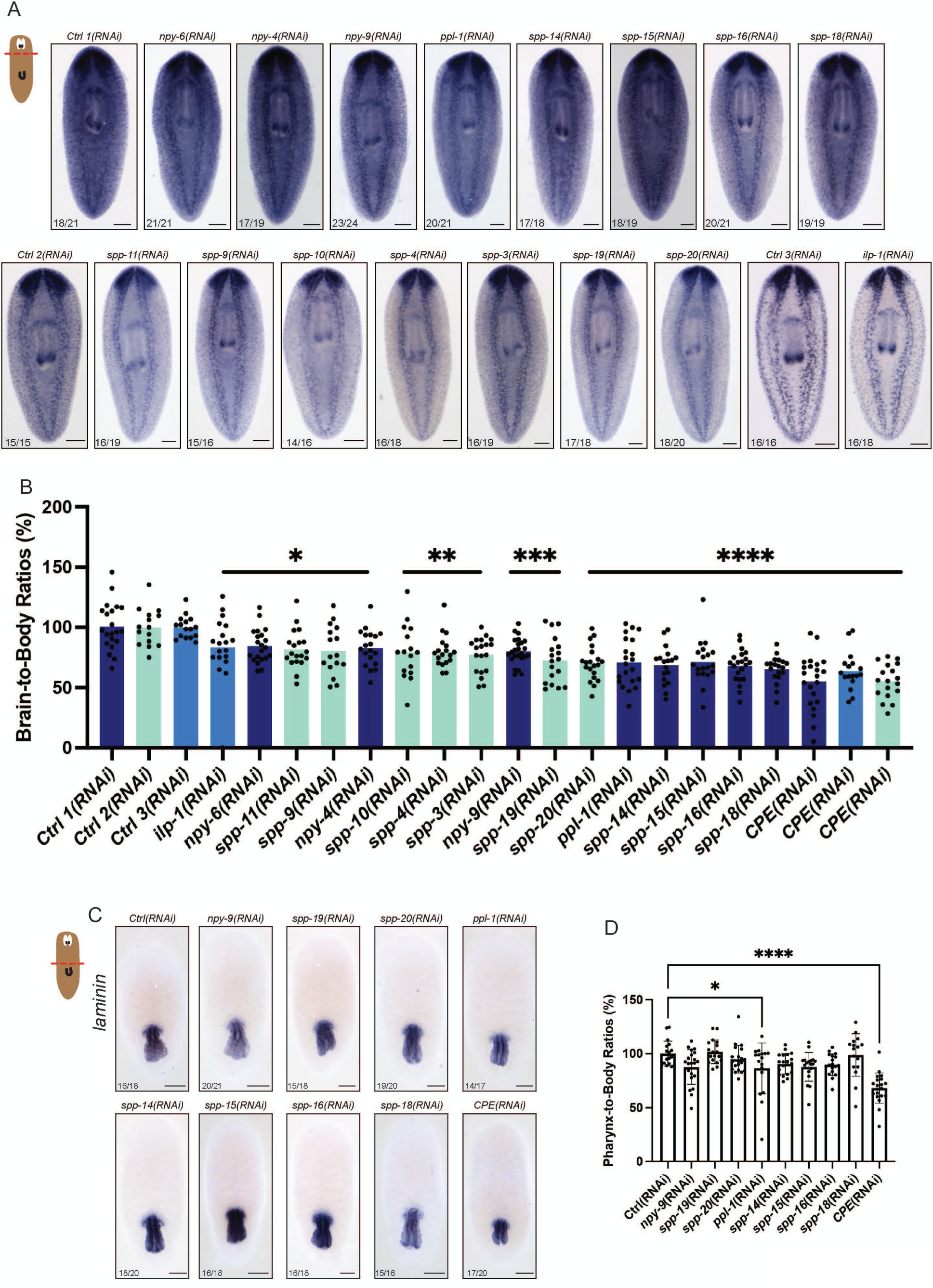
Neuropeptide-encoding genes promote robust brain regeneration. (A) Representative ISH images of all three RNAi groups which underwent the same RNAi paradigm of five 5 µg dsRNA feedings 3-4 days apart, head amputation, and killing and fixing 7 days after amputation. Worms were stained with *choline acetyltransferase*, a broad neuronal marker. (B) Graph showing brain-to-body ratios after RNAi and 7 days regeneration. Each RNAi group with its corresponding negative [*Ctrl/GFP*(RNAi)] and positive control [*CPE*(RNAi)] is represented in a different shade of blue/green. (N= 16-21 worms) (C) Representative images of heads regenerating tails and pharynges stained with *laminin*, a pharynx marker (Bohr et al., 2021). (D) Bar graph displaying pharynx-to-body ratios produced by tracing the pharynx and body in triplicate. (N=16-20 worms) Quantification for both A and C was completed with a One-way ANOVA and Dunnett’s multiple comparisons test. Both graphs display the means with standard deviation. * p < 0.05, *** p < 0.001, **** p ≤ 0.0001. Scale bars = 200 µm.

After identifying neuropeptides that promote regeneration, we wanted to determine whether their roles in regeneration occur via regulation of stem cells. First, we knocked down the eight neuropeptide-encoding genes with the most significant roles in regeneration and performed a head amputation. After seven days, we observed less *smedwi-1* mRNA expression after *spp-19*, *ppl-1*, and *spp-16* (Fig. 5A, indicated with arrowheads). To get a more quantitative look at this subtle phenotype, we also performed sublethal irradiation after RNAi targeting the same eight neuropeptide-encoding genes and observed the behavior of surviving individual stem cells. After RNAi targeting *spp-19*, *ppl-1*, *spp-14*, *spp-16*, or *spp-18*, animals had fewer total *smedwi-1^+^*stem cells per body area (Fig. 5B-D). Interestingly, we saw that RNAi of *spp-20* caused significantly fewer colonies and knockdown of *spp-19*, *spp-20*, *ppl-1*, *spp-14, spp-16* and spp-18 significantly reduced the number of stem cells per colony, indicating failure of surviving stem cells to divide appropriately (Fig. 5E-F). Interestingly, at least one gene that was required for regeneration—*spp-15*—impacted regeneration of the brain without impacting stem cells in our experiments, suggesting that neuropeptides play both stem cell-dependent and stem cell-independent roles in regeneration. Taken together, neuropeptide processors produce a diverse set of planarian neuropeptides that together promote the robustness of planarian regeneration through stem cell maintenance and function (Fig. 6A).

**Figure 5.**
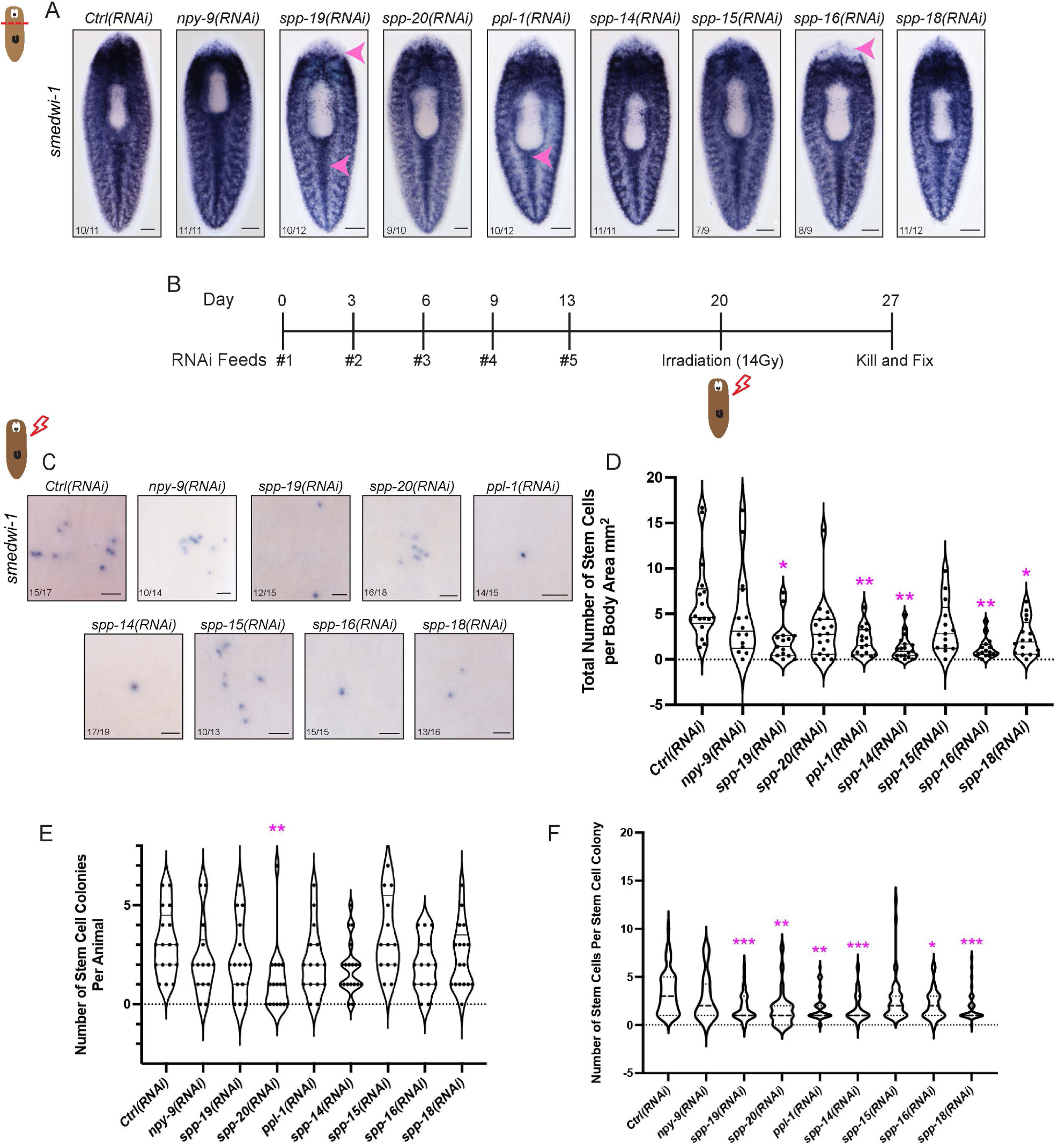
Neuropeptides promote stem cell survival and expansion. (A) Representative head regenerates after RNAi stained with *smedwi-1*, a stem cell marker. Magenta arrowheads indicate areas with less *smedwi-1* expression. The RNAi paradigm consisted of five 5 µg dsRNA feedings 3-4 days apart followed by a head amputation and 7 days to regenerate. (N= 9-12 worms) (B) The RNAi paradigm for sublethal irradiation experiments included five 5 µg dsRNA feedings 3-4 days apart, followed by sublethal irradiation at 14 Gy and 7 days of recovery post-irradiation before fixation for ISH. (N= 13-19 worms) (C) Insets of irradiated worms displaying *smedwi-1* mRNA expression. (D) A bar graph shows the total number of stem cells normalized to body area. Comparisons were completed with a Brown-Forsythe and Welch’s ANOVA test due to differences in variance. (E) We also quantified how many stem cell colonies were present in each animal (One-way ANOVA with Dunnett’s multiple comparisons test). (n=20-53 stem colonies counted per animal) (F) Finally, we quantified how many stem cells were present per stem cell colony (Brown-Forsythe and Welch’s ANOVA). * p < 0.05, ** p < 0.01 *** p < 0.001. Scale bars A = 200 µm and C = 50 µm.

**Figure 6.**
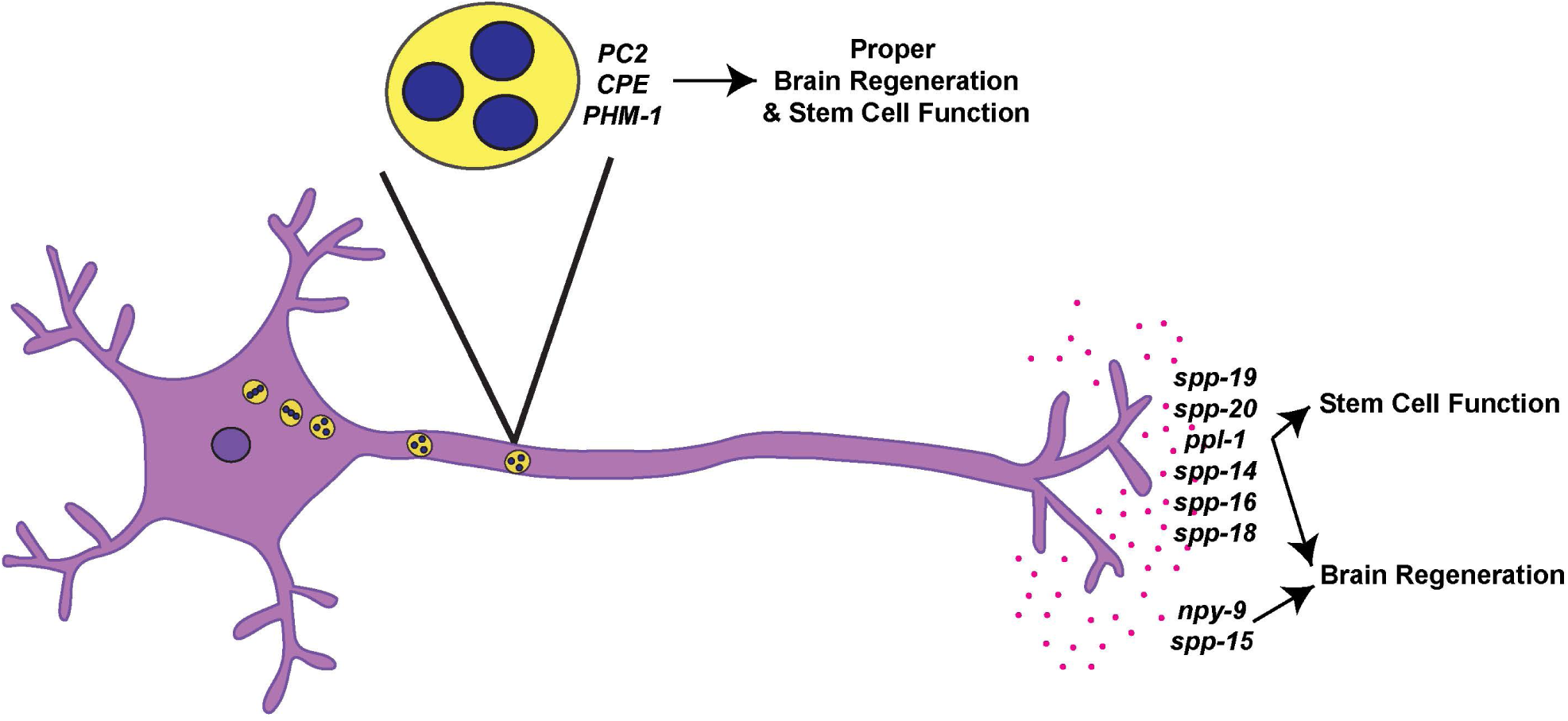
Summary diagram of neuropeptide processors and neuropeptide-encoding genes. A diagram of a neuron with the important neuropeptide processors for brain regeneration and the neuropeptide-encoding genes that impacted both brain regeneration and stem cell function.

## Discussion

We began this work seeking to determine whether the planarian nervous system promotes regeneration. Our work demonstrates a clear and convincing role for the nervous system in planarian regeneration and in pluripotent stem cell regulation. Further, we show that one essentialway the nervous system promotes regeneration is through complex neuropeptide signaling. We illustrated this essential role in two parallel ways. First, we identified three neuropeptide processing enzymes (PC2, CPE, and PHM-1) that collaborate to promote proper brain regeneration, stem cell maintenance, stem cell differentiation, and animal movement.

Second, to identify relevant targets of these enzymes, we queried the neuropeptide complement and found that RNAi of sixteen separate prepropeptide-encoding genes reduced the ability of planarians to regenerate. We confirmed that at least six of these neuropeptides impact regeneration *and* stem cell function. Our results suggest that neuropeptide signaling is necessary for stem cell function and successful regeneration in planarians. More broadly, our work indicates that the planarian nervous system itself is critical for regeneration, at least in part because it produces neuropeptide signals.

Though roles for most neuropeptides in our analysis are unknown, one of the neuropeptide hits from our regeneration study, *ilp-1,* encodes an insulin-like peptide with conserved roles in animal growth and metabolism across the animal kingdom (Chan & Steiner, 2000; Chowański et al., 2021; Fernandez & Torres-Alemán, 2012; Miller & Newmark, 2012). In planarians, RNAi of *ilp-1* caused animal degrowth, decreased mitotic stem cells, and reduced germline cells in sexual animals (Miller & Newmark, 2012). *PC2(RNAi)* and *CPE(RNAi)* animals seem to degrow, which could occur through loss of insulin signaling (ilp-1/inr-1) (Miller & Newmark, 2012). Another hit from our screen, pyrokinin peptide like (*ppl-1*), has been studied in other invertebrates. In cockroaches, a neuropeptide called leucopyrokinin which is part of the pyrokinin family, was found to be important for hindgut movement (Holman et al., 1986). Other pyrokinin-like neuropeptides have roles in feeding behavior (Bader et al., 2007). However, most neuropeptides in our analysis cannot be matched with neuropeptides in other organisms, either because they are planarian- or flatworm-specific or because their small size and divergence prevent detection of homology with conventional means. Further work will be required to determine whether some of these neuropeptides work through conserved mechanisms and whether these mechanisms in other organisms might be useful targets for pro-regenerative therapies.

Importantly, knock down of individual neuropeptide-encoding genes did not fully recapitulate phenotypes we observed after *PC2(RNAi).* One possibility is that combinations of planarian neuropeptides work cooperatively, which has been seen in other organisms. For example, in *C. elegans,* a knockout of three neuropeptides blocked stress-induced sleep; however, individual mutations of those same neuropeptides did not impact sleep (Nath et al., 2016). Importantly, we also acknowledge that additional neuropeptides might be encoded by the planarian genome, providing additional putative molecules to investigate. This possibility is supported by the detection of many uncharacterized, short transcripts expressed in peptidergic neurons in single-cell sequencing datasets (Fincher et al., 2018; Plass et al., 2018), many of which encode proteins with signal peptides and dibasic residues.

Based on our results, we believe that planarian stem cells are likely a direct or indirect recipient of neuropeptide signals. Thus, it will be important to determine the downstream signaling mechanisms activated by key neuropeptides. In the literature, neuropeptides most commonly activate G-protein coupled receptors (GPCRs) (Zhang et al., 2011) and over 550 GPCRs are encoded in the planarian genome, many of which are predicted to be neuropeptide receptors (Saberi et al., 2016). We previously characterized the heterotrimeric G-protein subunit complement in planarians, with an eye toward identifying factors that may act with key GPCRs in regeneration. We found that seven heterotrimeric G-protein subunits—six alpha subunits and one beta subunit—promote regeneration (Jenkins & Roberts-Galbraith, 2023); each is a candidate protein for mediating signaling downstream of key neuropeptides. Some of these G-proteins impact animal polarity and wound response, indicating that neuropeptides or other key neuron-derived signaling molecules may play key roles in these aspects of the wound response as well (Jenkins & Roberts-Galbraith, 2023).

Finally, one emerging role for neuropeptides is their ability to act as neuromodulatory signals to promote organism-wide states such as satiety or energy homeostasis (Deem et al., 2022; Krashes et al., 2016; van den Pol, 2012; Zhang et al., 2025). We hypothesize that a critical role of nervous systems in regeneration could be that neurons and/or neuroendocrine cells produce neuropeptides and other proteins, like newt anterior gradient protein, as a neuromodulatory mechanism to produce a pro-regenerative state necessary for activating and coordinating regenerative responses. In combination with literature on other organisms, our data show that this neuromodulatory function could be both a widely conserved element of regeneration and one that could be leveraged for development of regenerative therapies for humans in the future.

## Methods

### Animal Husbandry

Asexual planarians of the species *S. mediterranea* (CIW4 (Newmark & Sánchez Alvarado, 2000) were maintained at 18°C in 1X Montjuïc salts (1.6 mmol/L NaCl, 1 mmol/L CaCl_2_, 1 mmol/L MgSO_4_, 0.1 mmol/L MgCl_2_, 0.1 mmol/L KCl and 1.2 mmol/L NaHCO_3_ prepared in deionized or ultrapure water) (Cebrià & Newmark, 2005). Animals were fed regularly with locally sourced beef liver puree (White Oak Pastures, Bluffton, GA). For experiments, animals ranging from 1-5 mm were separated from stocks, starved, and treated with 1:1000 Gentamicin sulfate (50 mg/mL stock [Gemini Bio, West Sacramento, CA]) to reduce microbes in the water at least one week prior to starting the experiment.

### Identification of genes and cloning

Primers for desired genes of interest were designed using Primer3 or BatchPrimer3 to target ∼200-750 bp of the transcript. Genes were amplified and ligated into EAM5112-digested pJC53.2 vectors using standard molecular methods (Collins et al., 2010). A complete list of primers and plasmids can be found in (Supp. Table 2).

### Phylogenetic analysis and reciprocal BLAST

Potential neuropeptide processing enzymes were identified via BLAST of available transcriptomes (Fincher et al., 2018; King et al., 2024). We also identified homologous proteins from a variety of organisms (Supp. Table 3) and conducted phylogenetic analysis using the Phylogeny.fr platform (Dereeper et al., 2008). Alignment was conducted using MUSCLE (v3.8.31) with default settings and Gblocks to remove poorly aligned positions (v0.91b) (Edgar, 2022; Talavera & Castresana, 2007). The phylogenetic tree was constructed using PhyML (v3.1/3.0) with WAG replacement matrix (Guindon et al., 2010). Additionally, the tree was rendered using TreeDyn (v198.3). Reciprocal blasts were performed using NCBI BLASTx and we listed the top three or four matches from other organisms (Supp. Table 1).

### RNA interference (RNAi) experiments

After PCR amplification of constructs from plasmids, dsRNA was transcribed *in vitro* from the pJC53.2 vector containing a 300–1000 bp fragment of each gene of interest as described (Collins et al., 2010; Rouhana et al., 2013). dsRNA concentrations were estimated by gel electrophoresis of 1:10 dilutions with measurement of band intensity. For all neuropeptide processor RNAi experiments, 10-12 animals were transferred to 60 mm Petri dishes. For the neuropeptide-encoding gene RNAi experiments, 18-20 animals were used (20 worm separated into two 60 mm petri dishes). *PC2(RNAi)* experiments were performed through dsRNA feeding (Rouhana et al., 2013) once every three days for 2-3 total feedings (after which the animals became paralyzed and unable to eat). For each *PC2* dsRNA feeding, 3 µg dsRNA was mixed into ∼30 µL beef liver paste diluted 4:1 liver:salts. 1 µL green food dye was added to the mixture to visually confirm liver consumption. For the other neuropeptide processors (*PC1*, *PC-like*, *Furin-1*, *Furin-2*, *CPE*, *CPA/B*, *CPD*, *CPA/B/O*, *PHM-1*, *PAM-like*, *PAL-1)* and neuropeptide-encoding genes, 5 µg of dsRNA was fed 5 times with 3-4 days between feedings because these worms did not lose the ability to eat due to paralysis like *PC2(RNAi)* animals. dsRNA targeting *Aequorea victoria green fluorescent protein* (*GFP)* was fed as a negative control. For neuropeptide processor RNAi, *PC2(RNAi)* worms were used as a positive control and regularly lysed before killing and fixing. For all neuropeptide-encoding gene RNAis, *CPE*(RNAi) was used as the positive control. We chose to use *CPE*(RNAi) worms as a positive control because these worms did not lyse like *PC2*(RNAi) animals and we could complete the planned feeding schedule. After each feeding and periodically through experiments, animals were washed and put into fresh dishes and treated with 1:1000 gentamicin sulfate (50 mg/mL stock [Gemini Bio, West Sacramento, CA]). Images of live animals were captured using either an Apple iPhone SE or a Zeiss Axiocam 506 Color camera mounted on a Zeiss Axio ZoomV.16 microscope with ZEN 2.3 (blue edition) software (ZEISS Microscopy, Jena, Germany).

### Righting Assay

Worms were observed for up to five minutes after being placed on their dorsal side. Animals were timed for the number of seconds it took the worm to return to its ventral side (Jenkins & Roberts-Galbraith, 2023).

### Riboprobe synthesis & *in situ* hybridization (ISH)

Single-stranded, anti-sense riboprobes matching genes of interest were transcribed in the presence of a nucleotide derivative marked with Digoxygenin (Dig-11-UTP) (Roche) using standard molecular methods (Collins et al., 2010). ISH was performed as a modified version of the protocol previously described (King & Newmark, 2013). 7.5% or 10% N-Acetyl Cysteine (NAC) in 1x PBS (1.37 M NaCl, 27 mM KCl, 100 mM Na_2_HPO_4_, 20 mM KH_2_PO_4_, pH 7.4) was used as a reducing agent and anti-mucolytic. Additionally, we used 2 μg/mL Proteinase K solution instead of 5 μg/mL to prevent tissue damage. Hybridized riboprobes were detected with anti-digoxigenin Fab fragments conjugated with an alkaline phosphatase for subsequent development (1:2000 dilution) (Sigma Aldrich, St. Louis, MO). Post-antibody washes with TNTx (0.1 M Tris pH 7.5, 0.15 M NaCl, and 0.3% Triton X-100) occurred as follows: one 5-minute wash, one 10-minute wash, and six 20-minute washes. Colorimetric ISH staining development utilized nitro-blue tetrazolium (NBT) and 5-bromo-4-chloro-3’-indolyphosphate (BCIP) (both Roche) in an alkaline buffer (100 mM Tris pH 9.5, 100 mM NaCl, 50 mM MgCl_2_, 0.1% Tween- 20, brought up in 10% Polyvinyl alcohol) to promote phosphatase activity. Animals were cleared and mounted in 80% glycerol. Stained animals were imaged using a Zeiss Axiocam 506 Color camera mounted on a Zeiss Axio Zoom.V16 microscope running ZEN 2.3 (blue edition) Pro software (ZEISS Microscopy, Jena, Germany). All ISH images were taken on a PlanNeoFluar Z 1x/0.25 FWD 56mm objective with magnification ranges from 32x to 63x.

### Fluorescence *in situ* hybridization (FISH)

Probes for FISH experiments were made with either digoxigenin (DIG-11-UTP) or fluorescein isothiocyanate (FITC-12-UTP) (Roche). Probes were detected using anti-DIG-POD (Roche) or anti-FITC-POD (Roche). Experiments were conducted using the ISH protocol described above (King & Newmark, 2013). Few changes were made, including the development solution which included a TAMRA-tyramide solution [0.2% TAMRA-tyramide, 0.05% 4-IPBA, 0.05% H_2_O_2_ in TSA buffer (2M NaCl in 0.1 M Boric acid pH 8.5, filter sterilized) and quenching in sodium azide [100 mM sodium azide in PBSTx (PBS + 0.3% Triton-X100)] for 10 minutes to inactivate peroxidase. This was followed by additional blocking and antibody steps and development with FAM-tyramide (0.2% FAM-tyramide, 0.05% 4-IPBA, 0.05% H_2_O_2_ in TSA buffer). Samples were incubated with DAPI (10 µg/ml) (1:1000; Thermo Fisher Scientific, Waltham, MA) and washed. After one day, animals were mounted in VectaShield (Vector Labs) and imaged on a Zeiss 880 LSM confocal with Zen Black 2.3 SPI software under 40x magnification.

### Image Quantification and Statistical Analysis

To quantify brain regeneration, we generated brain-to-body ratios as previously described (Roberts-Galbraith et al., 2016). Triplicate brain and body area measurements were completed in FIJI (Fiji is just ImageJ) software (Schindelin et al., 2012). We normalized these values such that control animals were set to 100% for all experiments. In regeneration assays measuring blastema size, the same method was used to generate blastema/body ratios. Resulting data were graphed and statistically analyzed in Prism - GraphPad Version 9.0 software (GraphPad Software, San Diego, CA). Specific tests are included in the corresponding figure legends.

All RNAi experiments were blinded before quantification. RNAi experiments had at least 10 worms per knockdown condition. Negative controls (*green fluorescent protein*) and positive controls (*PC2*, *CPE*, or *hnrnpk*) were included for all RNAi experiments. For experiments with only two groups we used Welch’s unpaired t-tests as the statistical test. For all other experiments with >2 groups, a One-way ANOVA with Dunnett’s multiple comparison test was used.

### Quantitative Reverse Transcription Polymerase Chain Reaction (RT-qPCR)

To quantify transcript abundance, we extracted RNA from RNAi-treated animals with Trizol Reagent (Thermo Fisher Scientific, Waltham, MA) as per the manufacturer’s protocol. We degraded any remaining DNA in the samples with RQ1 RNase-free DNase (Promega Corporation, Madison, WI) at 37°C for a 15 min incubation. Purified RNA was used to synthesize cDNA using an iScript kit (Bio-Rad, Hercules, CA) as per the manufacturer’s recommendations. Reactions for RT-qPCR were completed with SYBR Green PCR Master Mix (Promega) in a QuantStudio® 3 real-time PCR system (Applied Biosystems, Foster City, CA). Primers targeting sequences ∼100 bp were designed with Primer3 (Koressaar & Remm, 2007; Untergasser et al., 2012), and primers meant to quantify RNAi target genes matched regions not included in the construct used for dsRNA. The gene *β-tubulin* was used to normalize for input in RT-qPCR reactions (1:2000 dilution) (Sigma Aldrich, St. Louis, MO). Triplicate biological replicates (12 animals each) were used for experiments, and RT-qPCR reactions were also completed in technical triplicate. Data were statistically analyzed and visualized using Prism - GraphPad Version 7.0 software (GraphPad Software, San Diego, CA). Specific tests employed are in the corresponding figure legends.

### Irradiation Experiments

For all sublethal irradiation experiments, worms were irradiated at 14 Gy 7 days after last RNAi feeding. The irradiator used was the CellRad+ non-medical irradiator from PRECISION x-ray irradiation. The total number of surviving stem cells was quantified using blinded files and all stem cells were individually counted using FIJI and normalized to the area of the body. Stem cell colonies were defined as clusters of *smedwi-1+* cells where no cells were >212 µm from the center of the colony. When the distance of 212 µm inches separated cells, we counted that as two separate colonies. We determined the 212 µm cut-off by evaluating each of the colonies in *gfp*(RNAi) worms and taking the average of all stem cell colony diameters.

## Supplement Figure Legends

**Supplement Figure 1.**
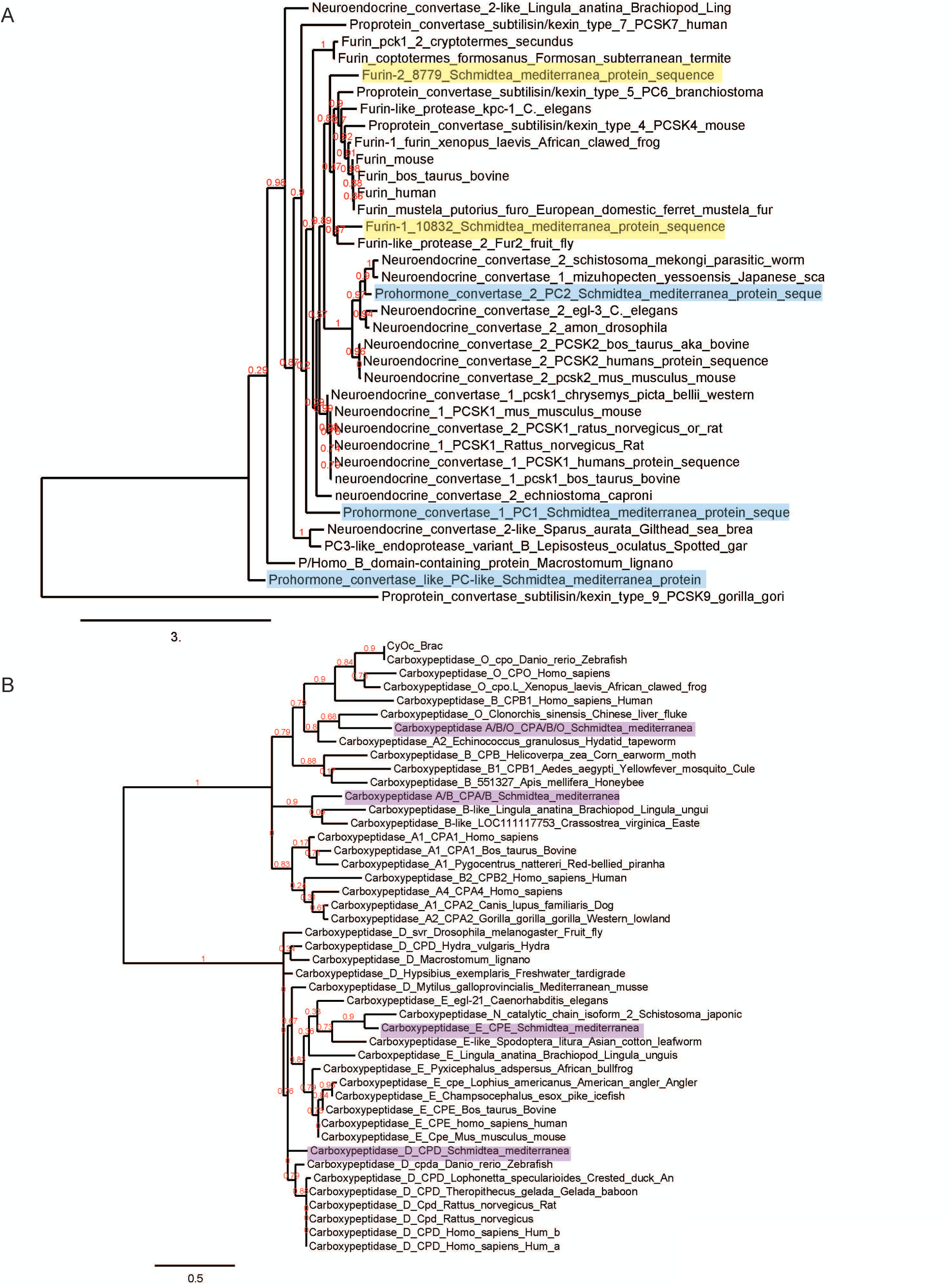
(A) Phylogenetic analysis of prohormone convertases and furins in other animals compared with those of the planarian, *Schmidtea mediterranea*. The key for the prohormone convertase tree is equivalent to 3 amino acid substitutions per site. The red numbers beside the branches indicate branch-support values. (B) The same parameters were used to construct the phylogenetic tree of carboxypeptidases. The planarian homologs for neuropeptide processors are highlighted. The key for this tree is equivalent to 0.5 amino acid substitutions.

**Supplement Figure 2.**
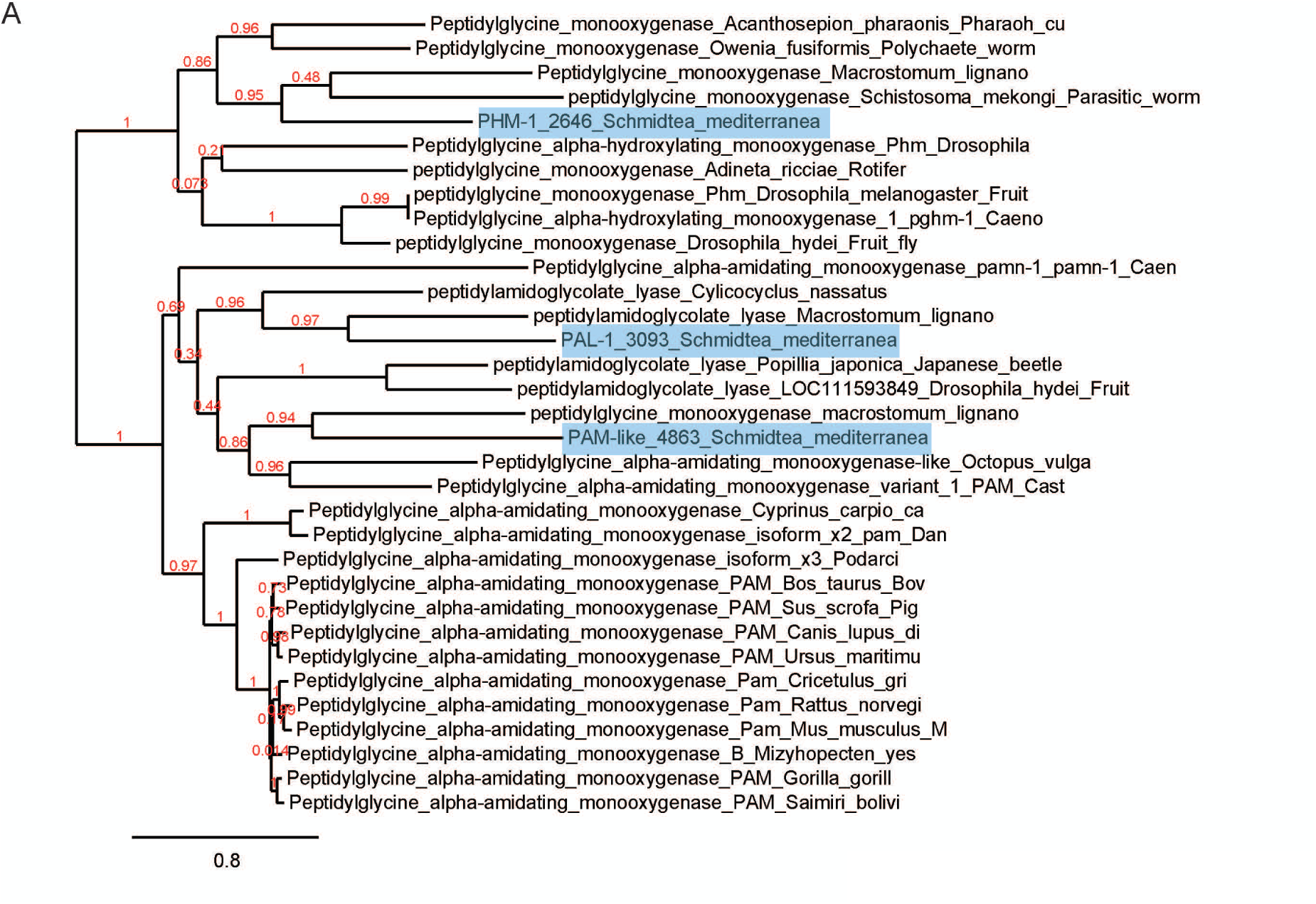
(A) Phylogenetic analysis of peptidylglycine alpha-amidating monooxygenases (PAMs) in various animals compared to prohormone convertases in planarian, *Schmidtea mediterranea*. The planarian homologs are highlighted in blue and the key is equivalent to 0.8 amino acid substitutions.

**Supplement Figure 3.**
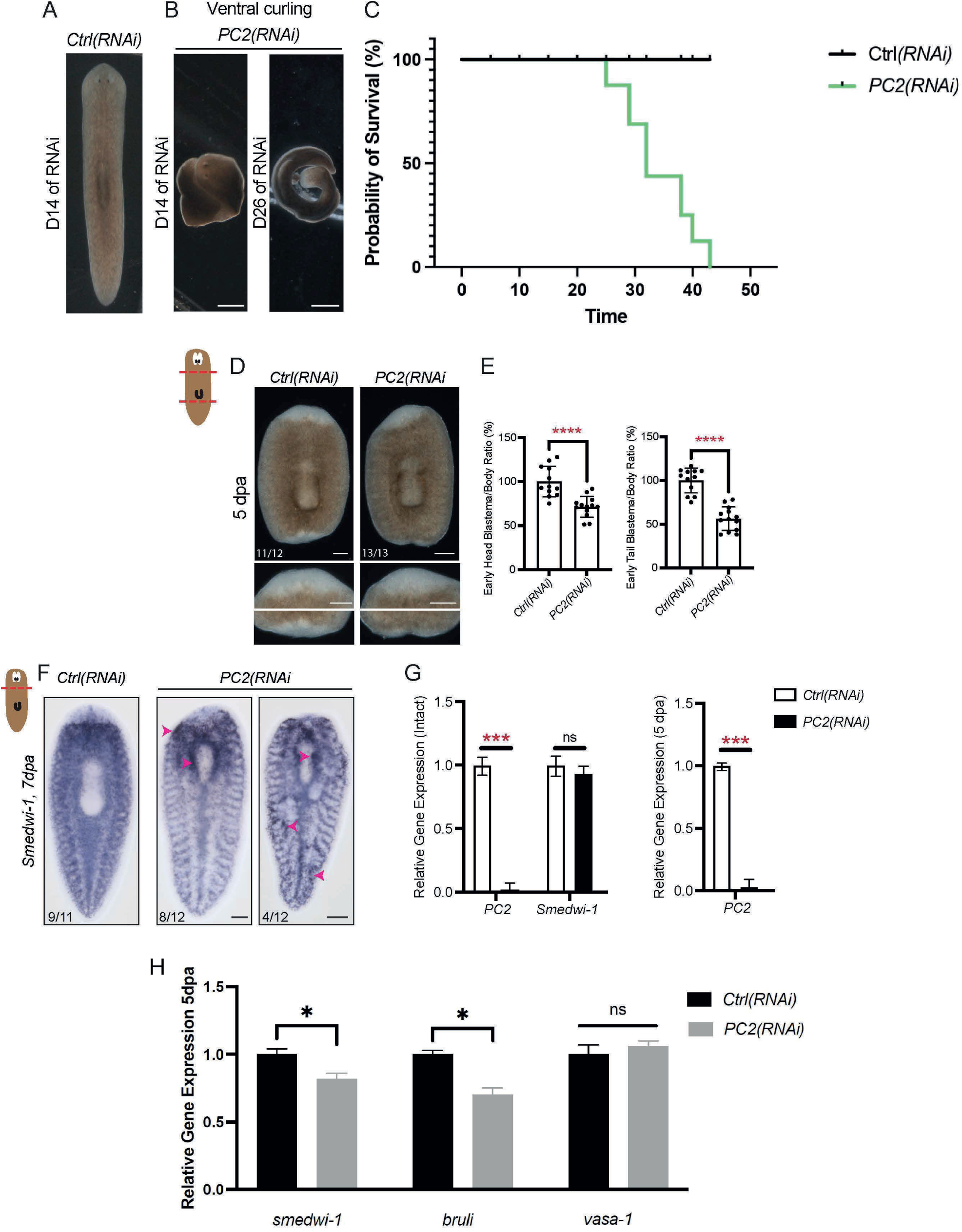
(A) Representative image of *Ctrl(RNAi)* fed *green fluorescent protein* dsRNA. (B) Image of ventral curling phenotype of *PC2(RNAi)* worms. (C) A survival curve shows the relative percentage of survival during RNAi. (N= 20 worms) (D) Representative images of head and tail regenerates after RNAi, 5 days post-amputation, and 13 days after the first RNAi feeding. Zoomed insets of the head and tail blastemas are included below the whole animal images. (E) A bar graph displays head and tail blastema/body ratios. Bar graphs are displayed as means with standard deviation. Data were statistically analyzed with unpaired t-tests. (F) *PC2(RNAi)* head regenerates stained with *smedwi-1* ISH. Magenta arrowheads indicate enriched stem cell expression near wound sites. (G) Relative expression levels of *PC2* in intact animals and 5 dpa animals and *smedwi-1* in intact animals, measured by RT-qPCR. (H) Relative mRNA expression levels of stem cell markers like *smedwi-1*, *bruli,* and *vasa-1* in 5 dpa worms. Error bars represent SEM. Differences in sample means were statistically analyzed with unpaired t-tests. * = p value ≤ 0.05, *** = p value ≤ 0.0005, **** = p value ≤ 0.0001. Scale bars B = 500 µm and D and G = 200 µm.

**Supplement Figure 4.**
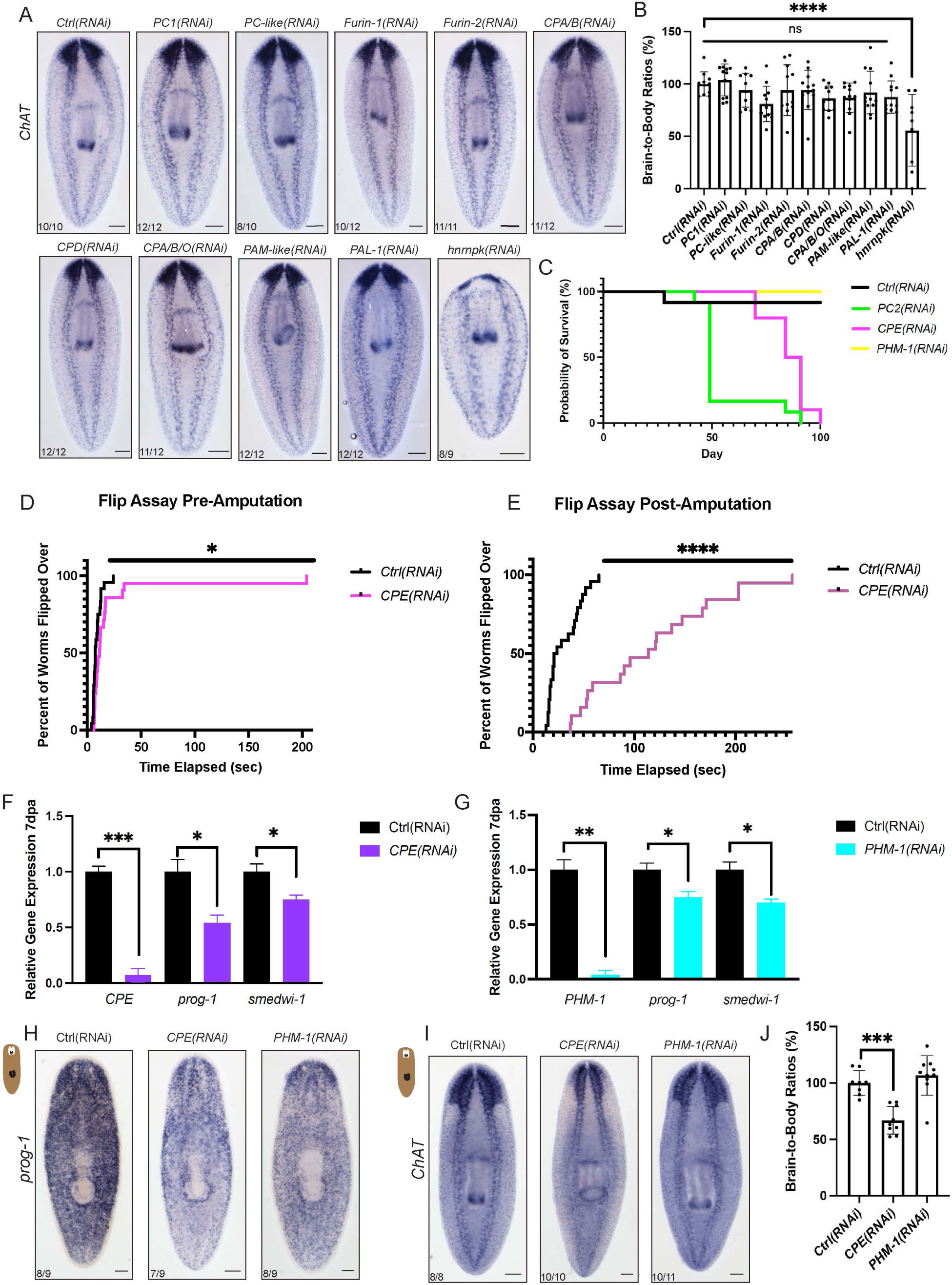
(A) Central nervous system expression shown by *choline acetyltransferase* (*ChAT*) ISH after RNAi of each neuropeptide processing enzyme-encoding gene. Positive control worms fed dsRNA for *heterogeneous nuclear ribonucleoprotein K* (*hnrnpk*). (B) Brain-to-body ratios for figure A completed with a One-way ANOVA and Dunnett’s multiple comparisons test (N= 9-12). (C) Survival curve shows the percentage of survival up to 100 days. One Ctrl(RNAi) worm died not because of the RNAi paradigm but because it crawled onto the lid of the Petri dish and dried out. (D-E) Graph indicating the percentage of RNAi-treated worms that flipped over after being placed ventral-side-up (D) pre-amputation (E) post-amputation. A Log rank (Mantel-Cox) test was used to compare results. (F) Relative expression levels of *CPE*, *prog-1*, and *smedwi-1* in amputated *CPE(RNAi)* worms were compared to *Ctrl/GFP(RNAi)* animals. (G) Relative expression levels of *PHM-1*, *prog-1*, and *smedwi-1* in amputated *PHM-1(RNAi)* worms compared to *Ctrl/GFP(RNAi)* animals. (H) Representative images of uninjured *CPE* and *PHM-1*(RNAi) worms stained with *prog-1*, an epidermal progenitor marker and (I) *choline acetyltransferase (ChAT),* a central nervous system marker. (J) Brain-to-body ratios are presented for non-injured RNAi animals. Quantification completed through a One-way ANOVA and Dunnett’s multiple comparisons test. Graph displays mean with standard deviation. * = p value ≤ 0.05, *** = p value, and **** = p value ≤ 0.0001.

**Supplement Figure 5.**
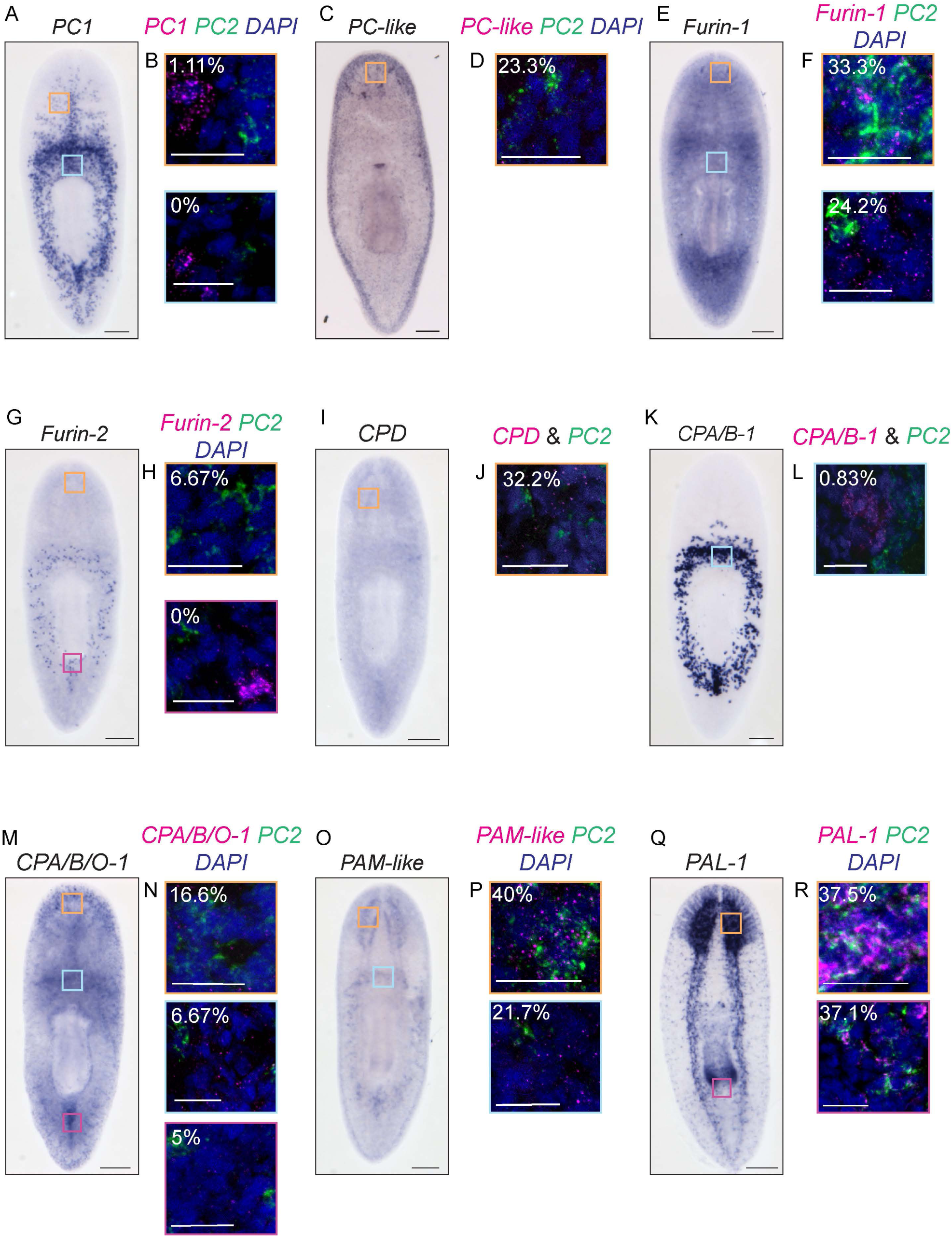
(A-R) Representative images of nine neuropeptide processing enzyme-encoding genes (*PC1*, *PC-like*, *Furin-1*, *Furin-2*, *CPD*, *CPA/B-1*, *CPA/B/O-1*, *PAM-like*, *PAL-1*) found in distinct locations across the body. For each neuropeptide processor, a colorimetric ISH is on the left with dFISH insets on the right showing the percentage of cells co-expressing *PC2.* (N= 6-7 worms; n= 60-240 cells counted per body region) For all insets, the gene of interest is in magenta, *PC2* is in green, and *DAPI* is in blue. Scale bars = 200 µm and inset scale bars = 20 µm.

**Supplement Figure 6.**
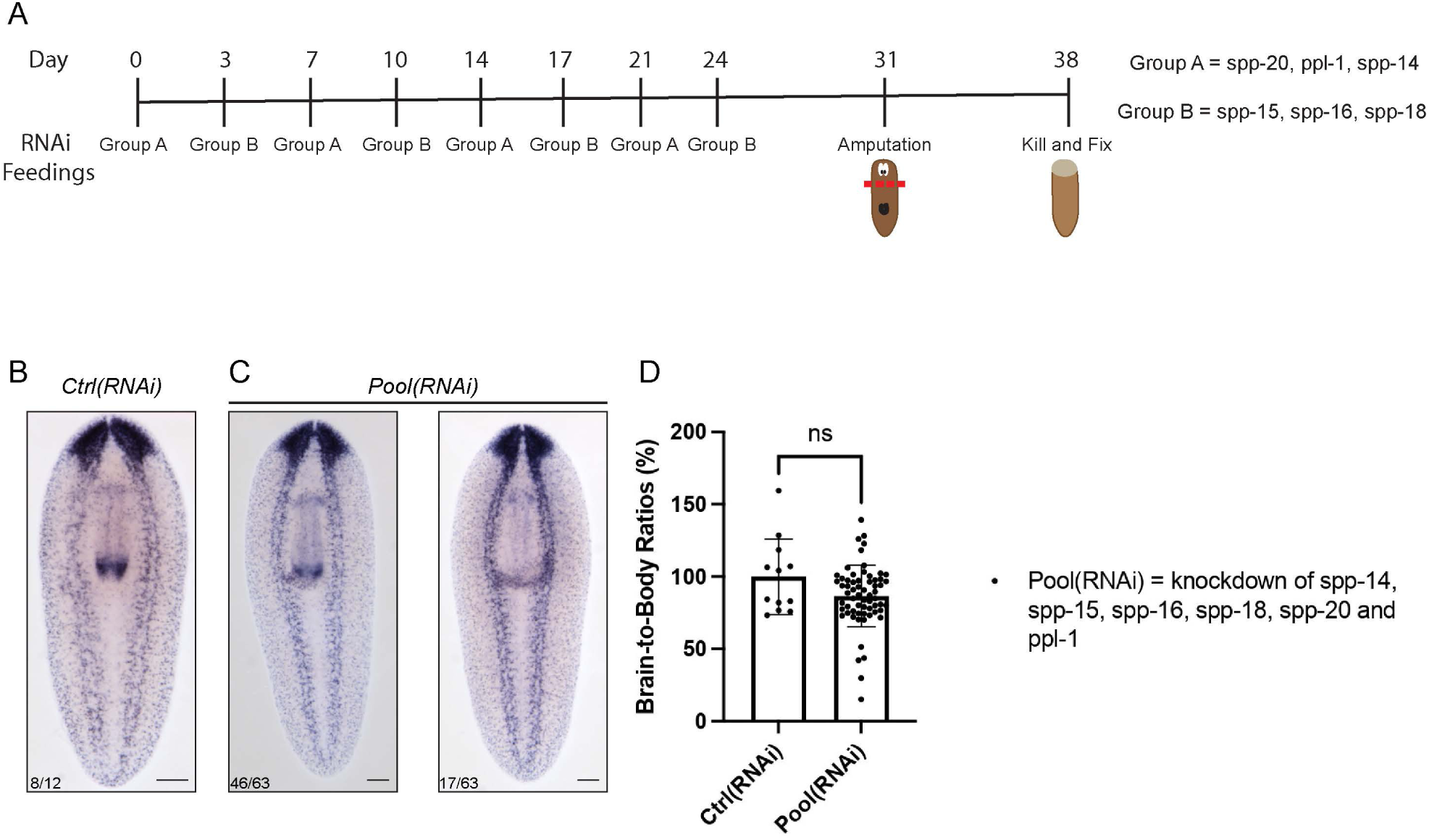
(A) Schematic of RNAi paradigm for multiple neuropeptide RNAi knockdown. Every 3-4 days, worms were fed 5 µg of dsRNA for either group A (*spp-20*, *ppl-1*, *spp-14*) or B (*spp-15*, *spp-16*, *spp-18*). Head amputation was performed seven days after the last feed and animals were allowed to regenerate for seven days. (B) Representative image of Ctrl(*RNAi)* worms fed *green fluorescent protein* (*gfp*) dsRNA. (C) *Pool(RNAi)* worms were fed dsRNA matching all six neuropeptides at least four times and these representative images display central nervous system expression through *ChAT*. (D) Bar graph showing brain-to-body ratios with means and standard deviation. Quantification completed through an unpaired t-test with Welch’s correction.

**Supplemental Table 1.** Reciprocal BLASTs for neuropeptide processors.

**Supplemental Table 2.** Plasmid and primer information for this study.

**Supplemental Table 3.** Sequences used in phylogenetic analysis.

## Conflict of interest statement

The authors declare no competing financial interests.

## Acknowledgments

We thank all previous and present Roberts-Galbraith lab members, especially Taylor Medlock- Lanier, Kendall Clay, Macey Wilson, Skylar Settles, Jazmine Dent, and Camina Gomes da Silva for support and comments on this manuscript. We thank Dr. Muthugapatti Kandasamy and the University of Georgia Biomedical Microscopy Core for help with fluorescent imaging. We thank Dr. Jim Collins, Dr. Phillip Newmark, and colleagues for generously providing neuropeptide plasmids. We thank the funding that allowed us to complete this work including the National Science Foundation CAREER award to RRG (IOS-1942822), National Institute of Health Genetics Training Grant (IOS-5T32GM142623) support of CEA, and the ARCS scholarship support for CEA. RRG is also supported by the Lois K. Miller Professorship through the Cellular Biology department at UGA. The earlier stages of this work were supported by a McKnight Scholars award and an Alfred P. Sloan fellowship to RRG. AEV was supported by a Pecot Fellowship from the McKnight Foundation and by funding from the Center for Undergraduate Research Opportunities at UGA, the Office of Experiential Learning at UGA, the McNair Scholars Program, and the Peach State Louis Stokes Alliance for Minority Participation which was funded by NSF grant 2207368.

## Notes

### Competing Interest Statement

The authors have declared no competing interest.

